# Twitchin kinase, a mechanoreceptor in the muscle sarcomere, is a catalytically-primed moonlighting kinase

**DOI:** 10.64898/2026.05.18.725903

**Authors:** Peter Gravenhorst, Frederic Berner, Hiroshi Qadota, Till Dorendorf, Tiankun Zhou, Rhys Williams, Michael Kovermann, Guy Benian, Olga Mayans

## Abstract

To explore conserved mechanisms and functions across mechanosensory kinases associated with the skeletal architectures of the cell, we investigated *in vitro* and *in vivo* the substrate targeting of twitchin kinase (TwcK), a mechanoreceptor from the muscle sarcomere. Specifically, we elucidated the crystal structure of TwcK in complex with substrates, used real-time ^31^P-NMR spectroscopy and luminescence-based assays to identify the phosphorylation site on a model peptide substrate, mined the *C. elegans* proteome to reveal the myosin regulatory protein MLC-4 as a substrate candidate and used CRISPR/Cas9 genome-edited and transgenic *C. elegans* strains to query the relation of twitchin and MLC-4 in muscle. Contrary to expectations, we find that TwcK undergoes activating conformational changes that are regulated by an N-terminal tail sequence that blocks hinge dynamics in the kinase fold. This distinct mechanism is conserved across sarcomeric, but not cytoskeletal, kinases. Functionally, cytoskeletal and sarcomeric kinases share an evolutionarily conserved phosphorylation targeting of myosin light chain (MLC) proteins. Yet, we find TwcK and its MLC4 substrate to segregate *in vivo* and not to constitute a functional kinase/substrate pair. Thus, canonical substrate targeting cannot be delivered by TwcK in its cellular context, where it has adopted a moonlighting role. We deduce this result to apply to other intrasarcomeric kinases. Our findings highlight how the cell context confers functional individuality to non-diffusible, otherwise conserved skeletal kinases.

## Introduction

Protein kinases are well-established chemosensors that mediate molecular communication in the cell in response to chemical cues. However, more recently, the phosphotransfer activity of several kinases has been linked to physical stimuli, especially mechanical force. Mechanosensing is particularly attributed to the family of titin kinases from the muscle sarcomere, which are regarded as *kinase mechanoreceptors*^1^. These conserved kinases are ubiquitously present in striated muscles across the animal biodiversity, being modules of giant, multi-domain filamentous proteins associated with the actin-myosin lattice of the sarcomere. Main representatives of this family are titin kinase (TK) in vertebrates, twitchin kinase (TwcK) and TTN-1 kinase (TTN-1K) in nematodes, and projectin kinase and stretchin-mlck in insects^1,2,3^. *In vivo*, these kinases experience pulling and shearing forces during muscle activity. Although their signalling pathways remain largely uncharacterized, mounting evidence suggests that they sense mechanical load on the sarcomere and translate it into biochemical events that inform other components of the cell of energy and physiological requirements, thereby contributing to trigger the adaptive response of muscle to mechanical demand^4,5,6,7^.

As in the specialized sarcomere, the actin-myosin lattice of the cytoskeleton is also contractile. It provides much of the force required for cellular activities such as motility, adhesion, cytokinesis and changes in cell morphology. Variations in the mechanical load of the cytoskeleton also trigger adaptive cellular responses, e.g. cell proliferation, differentiation and migration^8,9^. Interestingly, associated with cytoskeletal actin-myosin filaments are kinases that, evolutionarily, are closely related to sarcomeric kinases of the titin family. Leading representatives are death-associated protein kinases (<u>D</u>APK), myosin light chain kinases (<u>M</u>LCK) and the triple functional domain kinases (<u>T</u>rio). Jointly, sarcomeric and cytoskeletal kinases form the so-called DMT kinase clad^10,11^. Several cytoskeleton-associated kinases -including MLCK, DAPK and ZIPK/DAPK3-phosphorylate myosin light chain (MLC) proteins as cognate physiological substrates, some but not all in a Ca^2+^-calmodulin dependent fashion^12,13,14,15,16^. MLC proteins are key myosin regulators and their phosphorylation promotes actin-myosin contraction^17^. Notably, sarcomeric TwcK and TTN-1K robustly phosphorylate *in vitro* model substrates containing the MLC sequence targeted by cytoskeletal kinases^18,19,20,21,22,23^, raising the question of whether they also regulate actin-myosin motors in the sarcomere. Bringing support to this view, a genome-edited *C. elegans* strain that expressed an inactive mutated TwcK displayed a faster than normal swimming and crawling phenotype^6^, suggesting that TwcK activity impacts motor performance in muscle. In turn, force generation in the cytoskeleton has led to suspect that cytoskeletal members of the DMT clad have mechanosensory properties akin to those of sarcomeric kinases^10,24^. Taken together, the parallelisms have led to suspect shared mechanistic principles and functional roles across skeletal kinases at large.

The mechanisms by which mechanosensory kinases sense and translate force have been better studied in sarcomeric kinases. Crystal structures of TK^7,25,26^ and TwcK^22,27,28^ interpreted in the light of atomic force microscopy (AFM) data and steered molecular dynamic simulations (SMDS)^7,21, 22,29,30^ have led to propose a mechanism based on the elastic force-response of kinase accessory domains. Specifically, in TwcK and TK, the kinase catalytic core is flanked by N- and C-terminal tail extensions —termed N-terminal linker (NL) and C-terminal regulatory domain (CRD), respectively— that pack against the functional regions of the kinase fold (**Fig 1A,B)**. The NL tail binds against the hinge region, while the CRD tail docks onto the ATP- and substrate-binding sites, blocking the access of substrates to the active site. For *C. elegans* TwcK, it has been shown that the tails act synergistically to silence kinase activity^22^. AFM and SMDS data suggest that the pulling forces that emerge during sarcomere function cause the unravelling of the inhibitory tails from the kinase surface, resulting in the stretch-activation of phosphotransfer activity. Support to the physiological relevance of this mechanism has been brought by a study that monitored *in vivo* conformational changes in TwcK in the working muscle of freely moving *C. elegans* using FRET^31^. As cytoskeletal kinases also contain an inhibitory C-terminal tail extension, a similar mechanosensory mechanism is expected also for those kinases^24^.

**Fig 1:**
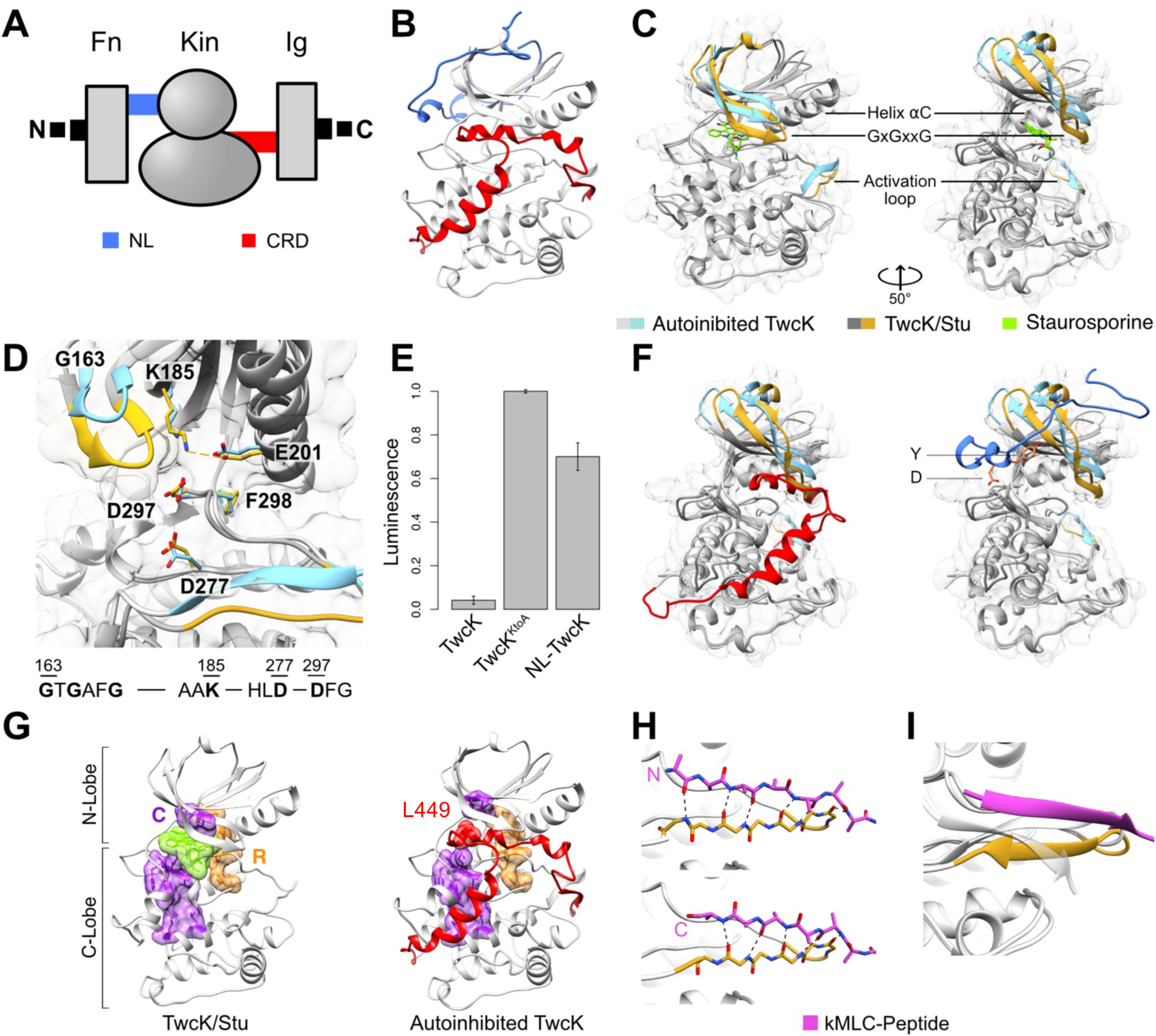
Structural characterization of TwcK activity states. **A.** Schematic domain composition of the kinase region of twitchin (Fn: fibronectin domain; Ig: immunoglobulin domain); **B.** Autoinhibited TwcK (extracted from PDB 3UTO^22^); **C.** TwcK (yellow) complexed to staurosporine (green) superimposed onto the kinase domain of the autoinhibited TwcK (PDB 3UTO) (cyan). In the latter, the flanking tail extensions have been removed to ease visualisation. Regions of the kinase fold retaining their conformation are shown in grey; coloured are the glycine rich β-hairpin and the activation loop, which are the functional elements displaying the largest shifts; **D.** Comparison of active site residues in the TwcK/Stu complex and in the autoinhibited TwcK. The sequence of catalytic motifs is shown below, with functional residues displayed in bold. For visual clarity, neither Stu nor the CRD are displayed in the corresponding states. The closure movement of the N-lobe towards the C-lobe in the Stu-bound TwcK leads to the formation of a salt bridge between the catalytically-essential residues K185 and E201 (indicated by a dashed line); **E.** Luciferase-based kinase activity assay. Data normalised to the negative control, inactive TwcK^K185A^, where the conserved lysine in strand β3 has been exchanged for alanine. Each histogram column is the average of three replicates (error bars are standard error). A reduction in luminescence signal correlates with kinase activity, so that signal absence corresponds to full consumption of ATP and, thus, maximal kinase activity. The NL-TwcK construction is largely inactive despite presenting a fully exposed and catalytically-primed active site; **F.** Superimposition of kinase states as in C. *Left:* the autoinhibited kinase (cyan) includes the C-terminal flanking tail (CRD; red). The CRD packs into the kinase ATP-binding pocket via its helix αR2 causing the glycine-rich β-hairpin to lift up. In TwcK/Stu (yellow), the glycine-rich loop swings down into a closed conformation that occupies the space previously filled by helix αR2. *Right:* The autoinhited kinase includes the N-terminal tail (NL; blue) packing against the kinase hinge region. The conserved YD motif is shown; **G.** Catalytic (C-) and regulatory (R-) spines are assembled in TwcK. The R-spine (orange) consists of residues: M205, L216, H275, F298; and the C-spine (purple) of residues: V170, A183, L238, M258, I283, M284, F285, L342, L346. The C-spine is completed by Stu in the active state or by residue L449 from the CRD in the autoinhibited state; **H.** Activation loop of TwcK complexed to the kMLC_13-22_ peptide (magenta) in parallel and antiparallel arrangement, as observed in non-crystallographic symmetry copies of TwcK. Dashed lines represent hydrogen bonds; **I.** Comparison of the peptide binding site of TwcK in its CRD-autoinhibited (PDB 3UTO; grey) and peptide-complexed states.

A central question in kinase mechanosensing is how the kinase can remain operational when placed under mechanical strain. In kinases, the active site is a large cavity located between the two lobes of the fold. The cavity accommodates the phosphate donor ATP/Mg^2+^ complex, while the peptide substrate docks onto the C-lobe^32^. For phosphotransfer to occur, the active site typically switches from an “off” to an “on” state via multiple and finely tuned conformational rearrangements that align catalytic groups into a productive state^32,33^. The changes are commonly triggered by the binding of ATP/Mg^2+^. How such elaborated changes can take place productively under mechanical strain is unknown. A current speculation is that, exceptionally, skeletal kinases do not require such inactive-active transitions, but instead they have a rigid fold locked into the active state that withstands mechanical deformation. This notion is based on crystal structures of autoinhibited TwcK and TK kinases that revealed the kinase core to be in a constitutive “on” state. Here, the inactive “off” state is achieved through the blockage of the active site by the flanking tails, which act as a *plug*^1,11^. Upon removal of this blockage, it is expected that the active kinase emerges. This view was further supported by 3D-structures of human DAPK1^34,35^, mouse DAPK2^36^ and the muscle pseudokinase PK1 from *Drosophila* obscurin^37^, which all showed apo states in catalytically productive conformations and no detectable structural changes upon ATP/ADP complexation. In summary, structural data leads to deduce that a rigid fold locked into the active state is the hallmark of skeletal kinases, which by bypassing the requirement of activation through fold dynamics can remain catalytically operational in their cellular contexts.

Despite the above described parallelisms, the existence of a shared operational principle across skeletal DMT kinases remains unclear. An impediment to address this question is that, to date, muscle kinases remain incompletely characterized. These kinases have only been observed in their tail-inhibited state and their cognate substrates are unknown. TwcK from *C. elegans* is the force responsive sarcomeric kinase best studied to date, having been characterized biochemically *in vitro*^18,20,22,23^, structurally^22,27,28^ and through mutagenesis *in vivo*^6,31^. In this study, we investigate a possible mechanistic and functional conservation across sarcomeric and cytoskeletal kinases by characterizing the active state of TwcK and its substrate targeting *in vitro* and *in vivo*. Contrary to expectations, our findings reveal mechanistic differences and highlight cellular compartmentation as a key mode of regulating the function of these non-diffusible kinases.

## Results

### TwcK undergoes activating conformational changes

To investigate the hypothesis of fold stiffness in kinases operating under tension, we characterised structurally the active state of TwcK from *C. elegans* by studying the isolated kinase domain without its flanking regulatory tails. Attempts to obtain a crystal structure of the uninhibited apo state as well as complexes with ATP/Mg^2+^, ATP analogues AMPPNP/Mg^2+^ or AMPPCP/Mg^2+^, or ADP/Mg^2+^ were unsuccessful. However, we succeeded in elucidating the crystal structure of TwcK in complex with staurosporine (Stu), a non-hydrolyzable ATP-analogue, to 2.40Å resolution **(Table 1)**. Crystals contained two molecular copies per asymmetric unit that were essentially identical (RMSD_Cα_=0.20Å across 268 atom pairs). Both TwcK molecular copies showed Stu bound in their ATP-binding pockets (**Fig 1B**; **Fig S1A**) and revealed that the removal of the CRD tail results in significant intramolecular flexibility at the unoccupied C-terminal lobe. In the ATP-binding pocket, the indole carbazole rings of Stu form extensive hydrophobic contacts with TwcK residues L162, G163, F167, V170, A183, K185, Y230, F232, E281, N282, M284, I296 as well as the aliphatic chain fraction of E237 and D297. Additionally, three hydrogen bonds are formed with the backbone of residues E231, M233 and E281. This binding mode is in agreement with that observed in other protein kinases, such as Stu-complexed DAPK (PDB: 1WVY) and PKA (PDB: 1STC).

**Table 1:** X-ray data statistics and model parameters.

|  | **TwcK/Stu** | **TwcK/Stu/kMLC<sub>13-22</sub>** |
| --- | --- | --- |
| PDB code | 29QN | 29UJ |
| Space group | P2<sub>1</sub> | P6<sub>1</sub> |
| Cell dimensions: |  |  |
| a,b,c (Å) | 71.79, 57.70, 86.09 | 109.13, 109.13, 105.52 |
| $\alpha, \beta, \mu$ (°) | 90, 105.42, 90 | 90, 90, 120 |
| Copies in a.u. | 2 | 2 |
| **Data Processing** |  |  |
| Beamline | PXII (SLS) | PXIII (SLS) |
| Detector | Eiger2 16M (Dectris) | Pilatus 2M-F |
| Wavelength (Å) | 1.000035 | 1.000 |
| Resolution (Å) | 47.4 - 2.40 (2.45 - 2.40)<sup>a</sup> | 48.16 - 2.50 (2.55 - 2.50)<sup>a</sup> |
| No. Reflections | 26696 (1556)<sup>a</sup> | 24765 (1415)<sup>a</sup> |
| R<sub>sym</sub>(I) (%) | 13.0 (141.2)<sup>a</sup> | 18.6 (317.4)<sup>a</sup> |
| < I/$\sigma$(I) > | 6.21 (1.07)<sup>a</sup> | 13.39 (0.94)<sup>a</sup> |
| CC1/2 (%) | 99.6 (55.8)<sup>a</sup> | 99.9 (47.8)<sup>a</sup> |
| Completeness (%) | 99.3 (99.9)<sup>a</sup> | 99.9 (99.9)<sup>a</sup> |
| Multiplicity | 3.68 (3.64)<sup>a</sup> | 14.02 (14.56)<sup>a</sup> |
| **Model Refinement** |  |  |
| No. working/free reflections | 26616/798 | 23840/864 |
| Rwork/Rfree (%) | 25.46/29.49 | 20.34/26.22 |
| No. Atoms Protein | 4295 | 4345 |
| No. Atoms Solvent | 136 | 174 |
| No. of Ligands / Atoms | 2 x Stu<sup>b</sup> / 70 | 2 x Stu<sup>b</sup> + 10aa peptide / 134 |
| Average B factors (Å<sup>2</sup>) |  |  |
| Protein | 93.13 | 74.63 |
| Solvent | 82.61 | 73.54 |
| Ligands | 77.90 | 75.09 |
| NCS R.m.s.d (Å) | 0.203 | 0.135 |
| R.m.s.d. bond length (Å) | 0.008 | 0.008 |
| R.m.s.d. angles (°) | 1.092 | 0.919 |
| Ramachandran plot |  |  |
| Favoured/Outliers (%) | 96.08/0.0 | 95.76/0.37 |

<sup>a</sup> Values in parentheses correspond to the highest resolution shell; <sup>b</sup> Stu: Staurosporine.

In canonical kinases, ATP binding induces the conformational transition of the kinase from an inactive to a catalytically committed state^32,33^. This transition involves *i)* the repositioning of helix αC (to allow for the formation of a catalytically-essential salt bridge between its conserved glutamate and the invariant lysine from strand β3, in the AxK motif, that coordinates the ⍺- and β-phosphates of ATP, productively orienting it for catalysis); *ii)* an open conformation of the activation loop to allow access to the active site; *iii)* the assembly of two spines (R[regulatory]-spine and C[catalytic]-spine) of stacked hydrophobic residues that promote the internal rearrangement of the kinase fold, including the mutual positioning of the lobes; and *iv)* the “in” arrangement of a conserved DFG motif, where the Mg^2+^-coordinating D residue is in the active cleft while the phenylalanine residue completes the R-spine^32,33^. In the autoinhibited TwcK, these features are already present, at the exception of *i)*. Accordingly, the occupation of the ATP-binding pocket in TwcK/Stu does not lead to global rearrangements and both the central DFG motif and helix αC retain their position. However, local conformational changes take place in the ATP-binding pocket and the peptide binding loop. In the ATP-binding pocket, the β-hairpin containing the glycine-rich motif is noticeably shifted in the TwcK/Stu complex, closing down onto the Stu molecule and locking it into the pocket (**Fig 1B**). The location of the glycine-rich loop in this closed conformation is occupied by helix αR2 of the CRD in the autoinhibited state; helix αR2 lifts the β-hairpin into a more open conformation (**Fig 1F**). In the TwcK/Stu complex, the invariant lysine residue (K185) in strand β3, which is immediately next to the moving β-hairpin in the same β-sheet, shifts to form a salt bridge with the conserved glutamate residue E201 from helix αC (distance change from 4.40Å ± 0.05Å in the autoinhibited TwcK in PDB 3UTO to 3.36Å ± 0.23Å in TwcK/Stu) (**Fig 1D**). A salt bridge between these two invariable residues characterises the catalytically committed state of kinases^32^. Thus, we conclude that the observed conformational change is activating.

To confirm that conformational changes at the ATP-binding pocket are a requirement for activity, we compared the phosphotransfer activity of the fully uninhibited TwcK with that of a NL-TwcK variant containing the N-terminal tail. The NL binds around the kinase N-lobe, onto the hinge region, and leaves the kinase active site fully accessible to substrates (**Fig 1F**; *right*)^22^. Luminiscence-based assays proved that the NL-TwcK variant has significantly less activity (∼65% reduction) than TwcK (**Fig 1E**), confirming earlier data^22^. The TwcK/Stu structure shows that, by wedging itself under the moving β-hairpin, the NL prevents its necessary motion. We conclude that substrate access to the active site is not sufficient for phosphotransfer catalysis but that conformational transitions are required for activity.

Further changes occur in the activation loop. In kinases, this loop serves as the binding platform for the peptide substrate. In the autoinhibited TwcK, this loop forms a β-strand that packs against the C-terminal strand βR1 from the CRD, forming a 2-stranded antiparallel β-sheet. The TwcK/Stu structure reveals that, upon removal of the CRD and thereby dissolution of the β-sheet, the activation loop undergoes a reorientation of its main chain, but retains an extended conformation and remains well-structured in electron density maps, being primed for substrate docking (**Fig 1B**).

### Investigation of the peptide binding site of TwcK

To understand the interaction of TwcK with its protein substrates, we studied next the peptide binding site using the 13aa long kMLC_11-23_ peptide model substrate^18,20,22,23^. While this did not yield diffracting crystals, a structure of TwcK in complex with Stu and the 10aa long peptide kMLC_13-22_ (TwcK/Stu/kMLC_13-22_) could be obtained to 2.5Å resolution (**Table 1; FigS2)**. Crystals contained two molecular copies of TwcK per asymmetric unit that were related by a two-fold and essentially identical (RMSD_Cα_=0.135Å across 268 atom pairs).

The conformation of TwcK in the TwcK/Stu/kMLC_13-22_ and TwcK/Stu complexes was closely similar (RMSD_Cα_=0.43Å), showing that peptide binding does not induce additional structural changes in TwcK. The Stu ligand was observed in the ATP-binding pocket as before, but the kMLC_13-22_ peptide lied astride two molecular copies of TwcK in the asymmetric unit. The peptide docked onto the activation loop (residues 164-QSVKV-168) of the two TwcK molecules, forming antiparallel and parallel β-strands respectively (**Fig 1H; Fig S2**). We deduce that the removal of the CRD-βR1 element is sufficient to induce the formation of a peptide binding site (**Fig 1I)**. Notably, the catalytic aspartate, D277, remained unoccupied and distant from candidate phosphorylation sites, leading us to speculate that Stu, which is larger than ATP and does not engage magnesium ions while binding to the kinase as ATP does, might not have induced all required rearrangements in the kinase.

### Pseudosubstrate roles of the C-terminal tail

In kinases, ATP binding leads to the formation of the C-spine by means of intercalation of the adenine base with the hydrophobic residues of the spine, which leads to the restructuring and stabilization of an active kinase conformation^32^. In TwcK/Stu, Stu completes the catalytic spine of TwcK (**Fig 1G**), but this does not result in lobe rearrangements as in canonical kinases. This is because in the CRD-autoinhibited TwcK (PDB 3UTO^22^) the two hydrophobic spines that traverse the kinase fold are already assembled and the DFG motif is in the “in” conformation (**Fig 1D,G**). A visual inspection of the autoinhibited TwcK reveals that this catalytically primed conformation is induced by the binding of the CRD into the ATP pocket. Here, residue L449 from the CRD helix αR2 is inserted into the spine, mimicking the role of adenine in ATP (**Fig 1G**). Nine out of the thirteen residues of TwcK that interact with Stu also form interactions with the CRD helix αR2. Thus, we conclude that helix αR2 is not a mere blocker of the ATP-binding pocket, but an ATP-mimic that stabilises the catalytically primed kinase state.

In the peptide binding site, the removal of the CRD-βR1 is sufficient to induce the formation of an accessible peptide binding site. Upon substrate binding, also a β-sheet is formed. We concluded that the CRD-βR1 element keeps the activation loop unavailable and prevents the substrate peptide to dock. Thus, causing the loop to switch between substrate accessible and inaccessible conformations (**Fig 1H,I**). Taken together, these data reveal the molecular basis of the CRD pseudosubstrate mechanism in TwcK, in which helix αR2 and strand βR1 are the primary inhibitory elements.

### Identification of the target phosphorylation site in the kMLC peptide substrate

MLC proteins are phosphorylated by cytoskeleton-associated kinases on a conserved sequence motif (T**<u>S</u>**NVF) located in an intrinsically disordered N-terminal sequence, where a serine residue (underlined) is the primary phosphorylation site^17^. This phosphorylation promotes the contraction of the actin-myosin cytoskeleton. The kMLC_11-23_ model peptide substrate of TwcK (11-KKRARAA**TSNVF**S-23) contains that consensus motif (bold). To investigate targeting parallelisms between TwcK and cytoskeletal kinases, we set to identify the phosphorylation site preference of TwcK on kMLC_11-23_. The peptide contains three potential phosphorylation sites: T18, S19 and S23. We asked which site(s) become phosphorylated by TwcK and in which ratio. To address this question, we monitored TwcK phosphorylation reaction using real-time ^31^P NMR spectroscopy (**Fig 2A,B**). Here, we observed that resonance signals derived from ATP (d, - 6.62ppm; d, −11.22ppm; t, −22.02ppm; where s: singlet; d: doublet; t:triplet) decreased, while concurrently five new resonance signals (s, 3.40ppm; s, 3.16ppm, s, 1.92ppm; d, −6.81ppm; d, - 10.86ppm) appeared. Comparison with reference spectra of pre-phosphorylated substrate peptides (kMLC ^pT18^, kMLC ^pS19^, kMLC ^pS23^) permitted to assign the resonance signal emerging at 3.16ppm to residue T18. (A ^31^P NMR based kinetic analysis of T18 phosphorylation is provided; **Fig S3**). Analysis of ATP, ADP and AMP at concentrations derived from the spectrum of the final activity measurement enabled the assignment of the resonance signal at 3.40 ppm to AMP and of the resonance signals at −6.81 and −10.86 ppm to ADP while the minor resonance signal at 1.92ppm corresponds to free phosphate (**Fig S4)**. Phosphorylation of sites S19 and S23 was not observed, suggesting that these residues do not act as relevant secondary sites, and revealing T18 as the main phosphorylation site in kMLC_11-23._

**Figure 2:**
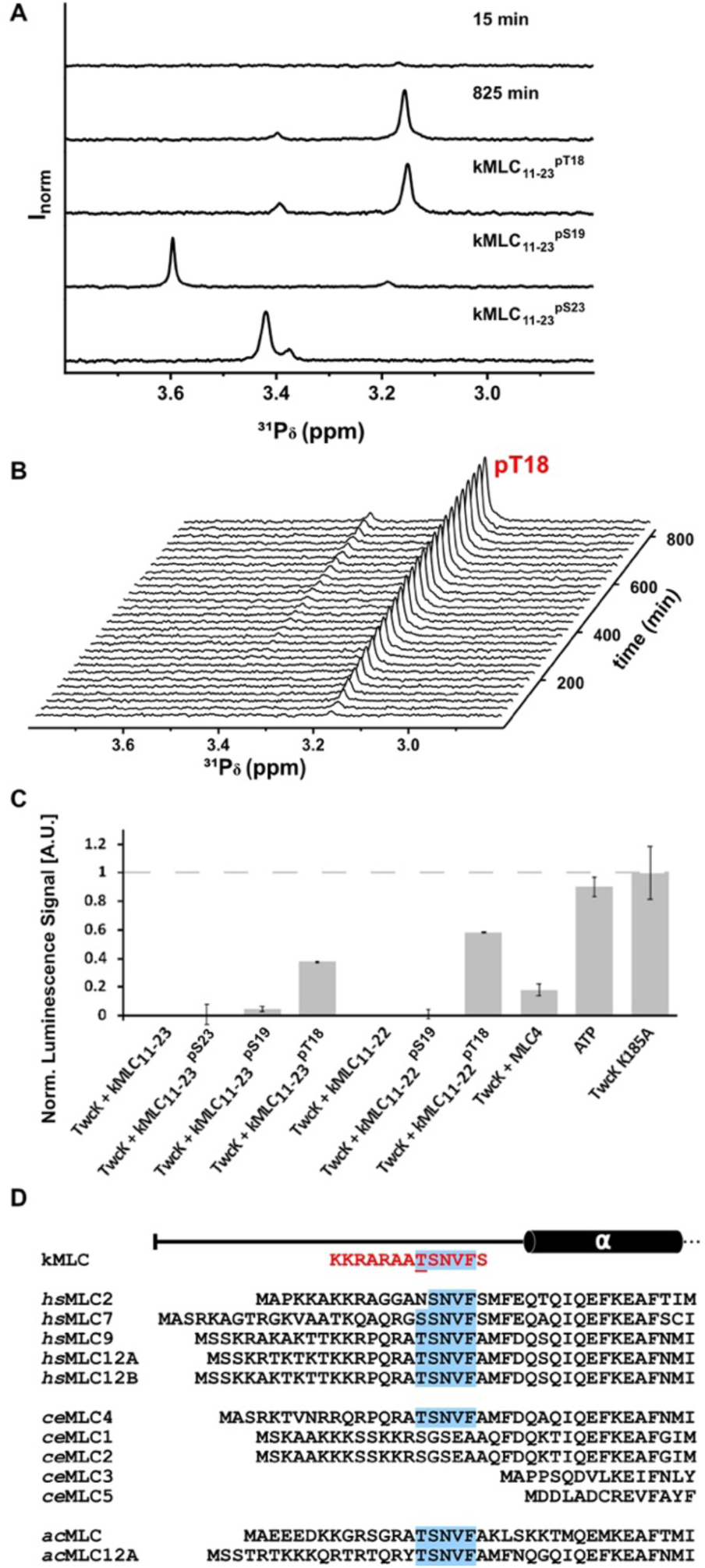
Monitoring TwcK phosphotransfer activity. **A.** 1D ^31^P NMR spectra of a reaction mixture containing TwcK, ATP, MgCl_2_ and kMLC_11-23_ plotted against incubation time. Only the spectral range of the emerging phosphorylated peptide product is shown; **B.** Spectral range 2.8-3.8 ppm of ^31^P NMR spectra shown in A. at the beginning of the reaction (i.e. after 15min incubation) and at the end of the reaction (825min incubation) compared to reference spectra of kMLC_11-23_^pT18^, kMLC_11-23_^pS19^, kMLC_11-23_^pS23^. ADP that is produced during the reaction decomposes over time into free phosphate (1.92ppm) and AMP (3.40ppm). Reference spectra for ATP, ADP and AMP are shown in **Fig S4**; **C.** Luciferase-based kinase activity assay. Data normalised to the negative control, inactive TwcK^K185A^, where the conserved lysine in β-strand β3 has been exchanged for alanine. Each histogram column is the average of three replicates (error bars are standard error). A reduction in luminescence signal correlates with kinase activity, so that signal absence corresponds to full consumption of ATP and, thus, maximal kinase activity; **D.** Sequence alignment of the peptide substrate model kMLC_11-23_ and N-terminal sequences of MLC proteins from *Homo sapiens* (*hs*) and *Aplysia californica* (*ac*) containing the conserved phosphorylation motif (highlighted in blue). For *C. elegans (ce)*, all MLC proteins are shown.

Results from NMR data were further confirmed by testing pre-phosphorylated peptide versions of kMLC_11-23_ in luminescence-based activity assays, which measure the ATP remaining in the medium upon termination of kinase activity **(Fig 2C)**. Here, a low luminescence signal is indicative of a high kinase activity. Both the pre-phosphorylated substrates kMLC_11-23_^pS19^ and kMLC_11-23_^pS23^ showed a luminescence signal comparably low to the unphosphorylated peptide, revealing that TwcK displayed full activity when the S19 and S23 phosphorylation sites were not available. To further confirm that S23 is not subject to TwcK phosphorylation, activity measurements on a kMLC_11-22_ peptide lacking this residue were performed that yielded a luminescence signal similar to that of kMLC_11-23_, showing that TwcK does not phosphorylate this terminal residue. The highest luminescent signal and consequently the lowest TwcK activity was consistently recorded for peptides kMLC_11-23_^pT18^ and kMLC_11-22_^pT18^, showing that once T18 is occupied by a phosphate group, kinase phosphotransfer is notably hampered. The remaining phosphotransfer activity recorded for kMLC_11-23_^pT18^ and kMLC_11-22_^pT18^ can be attributed to the known autophosphorylation of TwcK^18,23^. Taken together, the data showed that residue T18 is the single phosphorylation target of TwcK in kMLC_11-23_.

### *C. elegans* MLC-4 is an *in vitro* substrate of TwcK

While MLC was found as substrate of *Aplysia* TwcK in preparations of native mollusc tissue^19^, the kMLC_11-23_ peptide model substrate of *C. elegans* TwcK is a non-naturally occurring sequence developed by trialling a library of synthetic derivatives of chicken MLC^18,20^. Both *Aplysia* MLC and kMLC_11-23_ contain the conserved motif (TSNVF) of vertebrate MLC proteins (**Fig 2D**), but the best-established MLC proteins from *C. elegans* do not and are highly dissimilar to the synthetic kMLC_11-23_ (**Fig 2D**). In order to identify physiological candidate substrates of TwcK, we queried the *C. elegans* genome for the existence of proteins in this nematode containing the target phosphorylation motif. For this, we performed a sequence similarity search of the *C. elegans* translated genome using sequence RARAATSNVF contained in kMLC_11-23_ (see Methods). The protein sequence with the highest match score (21.4) to the search motif was myosin regulatory light chain isoform 4 (MLC-4) (RPQRATSNVF), a poorly characterized non-muscle specific MLC protein in *C. elegans*. No other prospective candidate could be identified in this way.

MLC-4 is the only MLC protein in *C. elegans* containing the canonical phosphorylatable motif of MLC proteins (**Fig 2D)**. To test whether MLC-4 is an efficient substrate of TwcK *in vitro*, we assayed a 13aa long peptide (RQRPQRATSNVFA) comprising its native sequence in luminescence-based phosphotransfer activity assays (**Fig 2C**). This showed that TwcK also displays strong levels of activity on the MLC-4 sequence, albeit slightly reduced (82%) compared to the synthetic kMLC_11-23_ peptide. A similar observation was made for *Aplysia* TwcK, where a peptide based on the native *Aplysia* MLC sequence was a worse substrate than the designer kMLC_11-23_^19^. Taken together, the data confirm that TwcK has robust levels of activity on the canonical MLC target motif, including that present in the endogenous MLC-4, displaying a substrate targeting typical of cytoskeleton-associated DMT kinases.

### *C. elegans* MLC-4 is an unlikely *in vivo* substrate of TwcK

MLC-4 in *C. elegans* is thought to be the only MLC protein in cytoskeletal myosin of non-muscle cells^38^. Although MLC-4 is not muscle specific, GExplore (v1.4)^39^ and SAGE data^40^ indicate that in *C. elegans mlc-4* mRNA is expressed at low levels in body wall muscle, pharyngeal muscle, and intestinal/rectal muscle. Conversely, it is well established that twitchin is robustly expressed in adult body wall muscle, where it localises to thick filament-containing A-bands^6,41^. Further, activity from *unc-22* (twitchin) promoters suggests the presence of twitchin also in pharyngeal and vulval muscles^6^. As TwcK is a domain of the non-diffusible twitchin protein, for MLC-4 to serve as its substrate the latter needs to associate with the sarcomere at the A-band. Currently, it is not known whether MLC-4 is present in the sarcomere. Thus, we explored here the localization of twitchin and MLC-4 in *C. elegans* muscle types where both proteins are jointly co-expressed; namely, pharyngeal and body wall muscle.

Using a *C. elegans* MLC-4-GFP expressing transgenic line (EU573, *orEx2*[mlc-4p::mlc-4::GFP; rol-6(su1006)^38^), we observed robust MLC-4 expression in the pharynx, but negligible expression in body wall muscle. When this nematode line was co-immunostained with antibodies to GFP to detect MLC-4-GFP and antibodies to MHC C (a myosin heavy chain isoform in the A-band of pharyngeal muscle), the signals did not overlap (**Fig 3A**). This indicated that MLC-4 was not recruited to sarcomeres in the pharynx. In fact, the cells that we observed to express MLC-4-GFP were pharyngeal gland cells^42^ and not pharyngeal muscle cells. In contrast, twitchin is expressed in pharyngeal muscle cells and integrated in sarcomeric thick filaments. The latter was validated by co-imaging the fluorescent signal from a CRISPR/Cas9 genome-edited *C. elegans* strain that expresses native twitchin tagged with mCitrine^31^ and antibodies to MHC C, which revealed a signal overlap confirming the sarcomeric A-band location of twitchin (**Fig 3B**). Lastly, we also examined the possible expression and localization of MLC-4-GFP in body wall muscle. Images of body wall muscle cells (**Fig 3C**), however, showed very weak MLC-4-GFP staining to small dots not localized in any pattern and not co-localizing with MHC B (a body wall muscle-specific myosin in the sarcomere). We concluded that MLC-4-GFP is not associated with sarcomeres in body wall muscle. These results suggested that in adult nematodes TwcK and MLC-4 are differentially compartmentalised.

**Figure 3.**
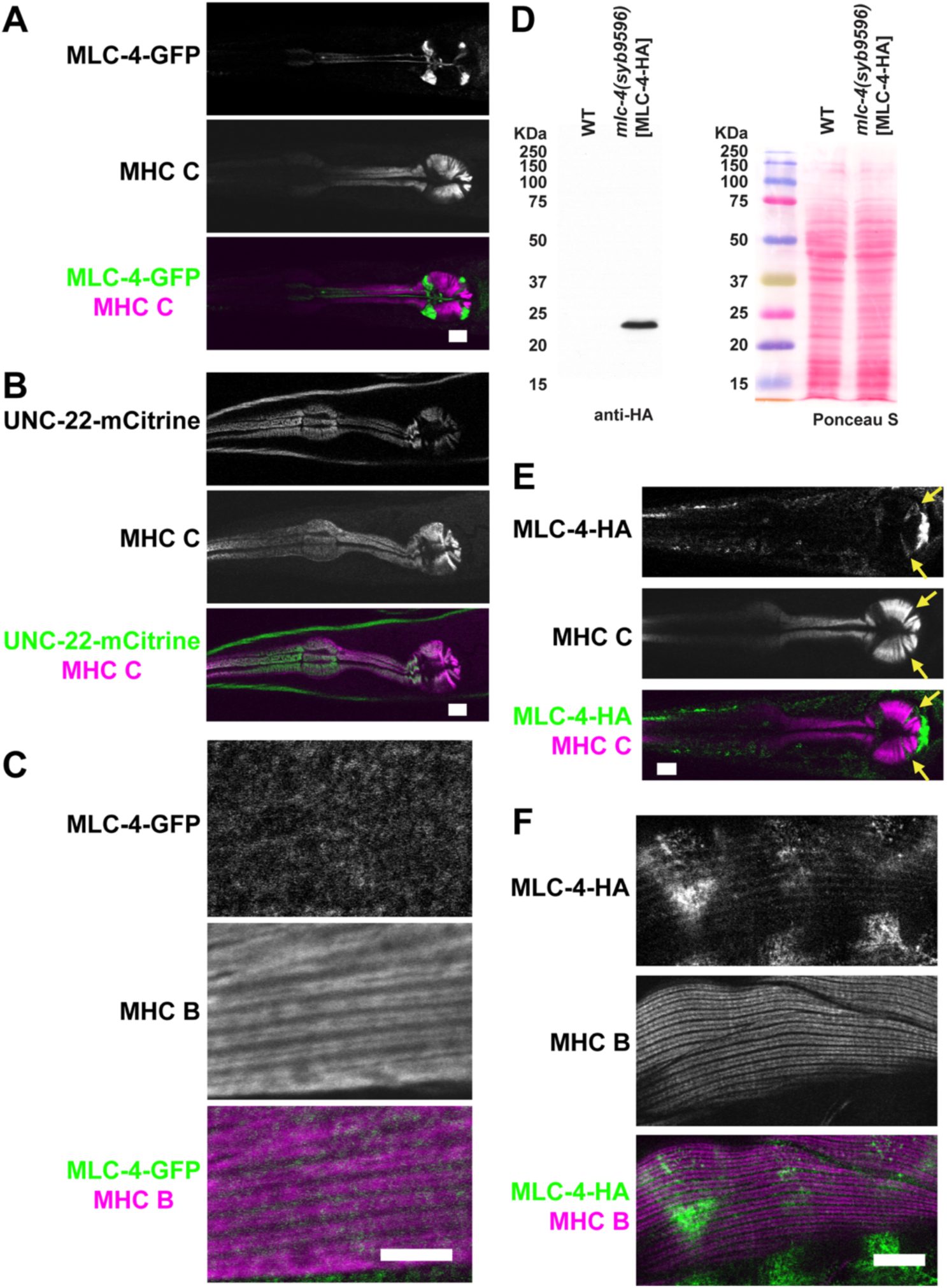
Twitchin and MLC-4 do not colocalize in *C. elegans* muscles. **A.** Confocal microscopy images of adult pharynxes from a transgenic line (EU573) expressing MLC-4-GFP co-stained with anti-GFP and anti-MHC C, a myosin heavy chain isoform expressed in pharyngeal muscle cells. MLC-4-GFP and MHC C signals do not overlap. MLC-4 seems to be expressed in pharyngeal gland cells; **B.** Images of the genome-edited strain PHX665 expressing twitchin tagged with mCitrine and immunostained with anti-MHC C. Twitchin is expressed in pharyngeal muscle cells, but not in pharyngeal gland cells. Note that twitchin is also expressed in body wall muscle (green signal at the periphery); **C**. Images of a portion of a body wall muscle cell from EU573 co-immunostained with anti-GFP and antibodies to the body wall muscle-specific protein myosin heavy chain MHC B in the sarcomeric A-band. MLC-4-GFP shows very weak staining as small dots not localized in any pattern and not co-localizing with MHC B; **D-F**. Expression of MLC-4-HA from its endogenous gene in the CRISPR strain PHX9596; **D.** A western blot reacted with anti-HA antibodies detects a protein of expected size from *mlc-4(syb9596)* [MLC-4-HA] and no detectable proteins from wild type (WT) worms; **E.** Co-immunostaining of *mlc-4(syb9596)* with antibodies to HA detecting MLC-4-HA and antibodies to MHC C that detect sarcomeric thick filaments in pharyngeal muscle cells. MLC-4-HA is present in pharyngeal gland cells (indicated by yellow arrows), but not pharyngeal muscle cells as shown by the lack of co-localization of MLC-4-HA and MHC C. The intense signal at the base of the pharyngeal terminal bulb may correspond to the pharyngeal-intestinal valve; **F.** Co-immunostaining of *mlc-4(syb9596)* with antibodies to HA to detect MLC-4-HA and antibodies to MHC B to detect thick filaments in the sarcomeric A-band of body wall muscle cells. There is no co-localization of MLC-4-HA and MHC B which suggests that MLC-4 is not localized to sarcomeric A-bands, the location of twitchin. The clusters of strong MLC4-HA signals likely arise from embryos or oocytes. (Scale bars, 10 μm).

To ensure that the apparent lack of expression of MLC4-GFP in body wall muscle from strain EU573 was not due to shortcomings of the transgene (e.g. due to the promoter region lacking the native gene 3’UTR that was not present), we generated a CRISPR/Cas9 genome-edited *C. elegans* line (PHX9596 [*mlc-4(syb9596)*]) expressing a HA-tagged MLC-4 protein produced from its own gene. Expression of MLC-4-HA in PHX9596 was validated by western blot using anti-HA (**Fig 3D**). Co-immunostaining with anti-HA and anti-MHC C revealed that, as with the transgenic line, endogenous MLC-4 is expressed in pharyngeal gland cells, not pharyngeal muscle (**Fig 3E**). Imaging of body wall muscle cells co-immunostained with anti-HA to detect MLC-4-HA and the body wall muscle specific anti-MHC B, revealed a weak presence of MLC-4 in body wall muscle –notably confirming the expression of the protein in this tissue–, but showed it associated weakly with the sarcomeric I-band (**Fig 3F**). The I-band localization of MLC-4 cannot be rationalized at this time, as no myosin is present in that zone. We speculate that this might be an artefactual association caused by the presence of the HA-tag in the protein. Importantly, the data reveal the absence of noticeable MLC-4 association with the sarcomeric A-band, where twitchin is located. These findings agree with earlier data^43^, where the major myosin heavy chains in body wall muscle, UNC-54 (MHC B) and MYO-3 (MHC A), were immunoprecipitated and the bound proteins identified using mass spectrometry. MLC-1, MLC-2 and MLC-3 were identified as myosin associated proteins in this way, but no MLC-4, hinting at MLC-4 not being present at appreciable levels in the sarcomeric A-band of body wall muscle. The non-sarcomeric location of MLC-4 is further supported by a study in which the ablation of MLC-4 expression using RNAi did not cause apparent defects in body wall muscle or pharyngeal muscle function in nematode embryos^38^.

Taken together, the data show that TwcK and MLC-4 are segregated in cells and that, even though MLC-4 contains the canonical phosphorylation target motif of DMT kinases, they do not form a functional kinase/substrate pair in muscle *in vivo*. Notably, the ablation of TwcK phosphorylation activity in a genome-edited nematode strain via a KtoA exchange that removes the catalytically essential lysine led to a motion phenotype^6^, suggesting that TwcK acts on another substrate(s) *in vivo*. In summary, our findings lead us to conclude that TwcK shares the evolutionarily conserved substrate targeting of cytoskeletal DMT kinases, but that this non-diffusible kinase is compartmentalized in a cellular context, where this canonical targeting cannot be delivered. We infer that in the sarcomere TwcK has adopted a moonlighting phosphorylating activity.

## Discussion

Kinases of the muscle sarcomere are regarded as paradigms of signalling mechanoreceptors, of which TwcK is the best studied representative. While stretch-activation has been proposed for these kinases^7,22,29,30,22,7^, fundamental mechanistic insights are lacking. In this regard, a basic requirement for phosphotransfer to be executed by a protein kinase is the correct structural alignment of its catalytic groups. Achieving such productive state requires the active centre to transition from an “off” to an “on” state through complex conformational changes, commonly induced by the binding of ATP/Mg^2+^ to the interlobular kinase region. A central question is whether the phosphotransfer activity of mechanosensory kinases displays a similar reliance on conformational dynamics. Data available to this date have led to think that skeletal kinases are locked into a constitutively active conformation, thereby evading the preparatory step of activation, with the “off” state achieved by the blockage of the active site by a terminal tail extension^1,11^. Contrary to expectations, the present data on TwcK show that removal of the CRD tail and occupation of the ATP-binding pocket induce activating changes, where closure of the glycine-rich β-hairpin leads to the formation of the catalytically essential K185-E201 salt bridge (**Fig 1**). Thus, contrary to cytoskeletal kinases^34,35,36^, TwcK requires activating conformational changes.

Remarkably, the restraint of the observed motion by the NL tail is sufficient to yield TwcK largely inactive, even when the active site is fully exposed (**Fig 1E**). This reveals the NL as an important intrasteric inhibitor of TwcK. SMDS have suggested that the NL tail is the primary mechanosensory element of TwcK, responding to stretch first^22^. Within the NL, a strictly conserved YD sequence motif has been recently identified across sarcomeric kinases, including TwcK and TTN-1K from nematode, insect projectin kinase and vertebrate titin kinase; kinases which otherwise comprise highly divergent NL sequences^26^. The motif anchors the NL tail to the hinge region in the kinase fold (**Fig 1F**;*Right*) and, according to SMDS, acts as a force barrier preventing the unravelling of the NL tail by stretch^7,22^. Results from the current study suggest that the YD motif plays a pivotal role in TwcK intrasteric inhibition by blocking the motion of the ATP-binding β-hairpin. Thus, it could be deduced that the motif serves as a *force-gating mechanism* to dictate the force at which the NL response ensues and, thereby, kinase activation. We conclude that the NL tail and its YD motif are key elements of a mechanosensory mechanism that is distinct and conserved in sarcomeric kinases.

Functionally, TwcK is most closely related to MLCK (51.6% seq.id.; **Fig S5**), that stimulates actin-myosin contraction via the phosphorylation of MLC^17^. MLC is also targeted by DAPK and ZIP3^12,14,15,16^. Phosphorylation chiefly occurs at residue S19, with T18 being a secondary site. TwcK from both *C. elegans* and *Aplysia*^19,20,22,23^ as well as TTN-1K^21^ also target the MLC motif *in vitro*. Thus, MLC could be regarded as a consensus substrate of DMT kinases. Our data show that T18 is the primary phosphorylation target of *C. elegans* TwcK and that S19 is not a significant secondary site. The reason for this differential targeting might reside on the location of basic residues in the kMLC_11-23_ peptide. An arginine residue is located in position 16 in native MLC sequences (**Fig 2D**). The shift of R16 to position 15 in non-natural peptides, such as kMLC_11-23_, resulted in MLCK completely switching its specificity for S19 to T18 (kMLC_11-23_ is peptide 11 in ^44^). Predictably, the observed targeting of TwcK in our study might also be the consequence of a specificity switch. This further highlights the close conservation in substrate targeting properties between TwcK and DMT kinases.

In *C. elegans*, only the protein MLC-4 contains the conserved MLC motif. We show that MLC-4 is an efficient *in vitro* substrate of TwcK (**Fig 2C**), while MLC proteins from *C. elegans* lacking this motif have been shown to barely elicit activity^20^. Despite this fact, our investigation of *in vivo* localization reveals that MLC-4 is an unlikely physiological substrate of TwcK. Twitchin is expressed in body wall muscle^41^ and pharyngeal muscle^31^, being present in thick filaments in A-bands. However, our data show that MLC-4 is not expressed in pharyngeal muscle and only lowly expressed in body wall muscle, but not localizing to the sarcomeric A-band (**Fig 3**). Thus, in adult muscle, TwcK and MLC-4 are non-diffusible proteins that do not co-localize. TwcK and MLC-4 are also not likely to interact during embryonic development because twitchin mRNA is not expressed in embryonic muscle^40^. In agreement, twitchin has not been detected in embryonic muscle by immunostaining and twitchin null worm mutants do not have any effect on embryogenesis^5,41^. Supporting the finding that MLC-4 does not associate with the sarcomeric A-band, immunoprecipitation of UNC-54 (myosin B) and separately MYO-3 (myosin A) from a mixed population of worms that included embryos, larvae and adults (myosin B and A are expressed in embryonic, larval and adult muscle) identified MLC-1, -2 and -3 as bound to myosin, but MLC-4 was not detected^43^. We thus conclude that TwcK and MLC-4 are spatially and temporally segregated *in vivo* and, thereby, are an unlikely kinase/substrate pair.

In summary, we infer that sarcomeric and cytoskeletal kinases of the DMT clad share an evolutionarily conserved substrate targeting pattern, but that sarcomeric kinases have evolved a distinct mechanosensory mechanism and are placed in cellular contexts where they do not necessarily act on canonical phosphorylation substrates. Since loss of TwcK activity in genome-edited nematodes carrying a KtoA dead-kinase point mutant has been shown to result in a fast swimming and crawling phenotype characterized by increased rates of muscle contraction/relaxation^6^, it must be deduced that in its cellular context TwcK acts on alternative substrates with atypical target sequences. We predict that these findings apply to other sarcomeric kinases, also shown to act on the conserved MLC sequence motif but with similarly discordant intrasarcomeric locations.

## Methods

### Cloning

The expression construct for TwcK from *C. elegans* comprising the catalytic domain (TwcK) (UniprotKB Q23551; residues 6251-6535) has been previously reported^23^.

To facilitate the direct comparison of crystal structures in this study and that previously reported for the multi-domain Fn-NL-TwcK-CRD-Ig kinase segment from twitchin (PDB entry 3UTO^22^), residue V6108 in UniprotKB Q23551 was taken here as residue number 1.

### Protein expression and purification

Protein production was as previously described^45^. In brief, protein overexpression was in *E. coli* Rosetta (DE3) cells (Merck Millipore) grown in Luria-Bertani medium and induced at an OD_600_ of 0.6-0.8 with 0.5mM isopropyl β-D-1-thiogalactopyranoside (IPTG) followed by incubation at 18°C for 18h. Cell lysis was by sonication in a solution of 50mM Tris-HCl, 500mM NaCl, 1mM DTT, pH 7.9 containing 20µg/ml DNase I (Sigma Aldrich) and cOmplete™ EDTA-free Protease Inhibitor Cocktail (Roche). Lysates were clarified by centrifugation and syringe filtered (0.22µm). Proteins were isolated from the supernatant by using Ni^2+^-affinity chromatography on a Histrap HP 5mL column (GE healthcare). Elution of His_6_-tagged protein was by continuous imidazole gradient and eluted samples exchanged into 50mM Tris-HCl, 50mM NaCl, 1mM DTT, 2mM MnCL_2_, pH 7.9 using either a PD-10 desalting column (GE Healthcare) or dialysis. To obtain the fully active dephosphorylated state^23^, TwcK samples were incubated with 400 units of Lambda protein phosphatase (NEB) in the mentioned buffer concurrently in the presence of TEV protease for His_6_-Tag removal (*ca* 18h, 4°C). Subtractive Ni^2+^-affinity purification was then performed on a 5mL HisTrap HP column connected to an Åkta FPLC system (GE Healthcare). Size exclusion chromatography used a Superdex 200 16/60 column (GE Healthcare) in 50mM Tris-HCl, 50mM NaCl, 0.5mM Tris(2-carboxyethyl)phosphine hydrochloride (TCEP), pH 7.9. Sample purity was assessed using SDS-PAGE.

### Structural elucidation of TwcK in complex with staurosporine

The TwcK sample was concentrated to 11.3mg/mL based on A_280_ and staurosporine (Stu) (Thermo Fisher) added at a final molar ratio of 1:2 (protein:staurosporine). The final concentration of TwcK upon addition of Stu was 10.5mg/ml. The mixture was incubated for 1h on ice before the setting of crystallisation trials. Crystallisation trials were set up in 96-well Intelliplates (Art Robbins) using the sitting drop, vapour diffusion method. Crystallisation drops consisted of 200nL TwcK/Stu mixture and 200nL reservoir solution. Trials were incubated at 18°C. Crystals of TwcK/Stu grew from 0.2M lithium sulphate, 0.1M bis-Tris pH 5.5, 25% [w/v] PEG 3350. For X-ray data collection, crystals were cryoprotected in mother liquor supplemented with 30% [v/v] ethylene glycol, prior to flash cooling in liquid N_2_.

X-ray data collection was performed on beamline PXII of the Swiss Light Source (SLS, Villigen, CH). Data processing used XDS/XSCALE^46^. Molecular replacement was performed with Phaser^47^. To account for possible interdomain motions, the N- and C-terminal lobe domains of *C. elegans* TwcK (extracted from PDB entry 3UTO and excluding flanking domains and regulatory tails) were used separately as independent search models. The molecular replacement solution was then refined in phenix.refine^48^ applying TLS refinement and non-crystallographic symmetry (NCS) restraints (each molecular copy of TwcK/Stu in the asymmetric unit was declared as a NCS unit, while each N- and C-lobe formed a TLS group). Manual model building was performed in Coot^49^. The Stu molecule was obtained from PDB 1AQ1. Statistics for X-ray data processing and model refinement are given in **Table 1**. The resulting TwcK/Stu model includes all protein residues at exception of 18 C-terminal residues, which were not visible in electron density maps.

### Structural elucidation of TwcK in complex with staurosporine and peptide substrate

Crystallization of TwcK in complex with Stu and a 10-aa long model peptide substrate (kMLC_13-22_; sequence RARAATSNVF) was performed by mixing 2.7mM kMLC_13-22_, 450µM Stu and 300µM TwcK in 20mM HEPES pH 7.6, 100mM NaCl. The final concentration of TwcK in the mixture was 8.6mg/mL. The peptide substrate was derived from a previously described 13-aa long sequence that had been diversified from native myosin light chain (MLC) from chicken and that proved to be an efficient *in vitro* substrate for TwcK^20,22^. The TwcK/Stu/kMLC_13-22_ complex was crystallized as described above from 2.5M ammonium sulfate, 0.1M bis-Tris propane pH 7.0. For X-ray data collection, crystals were vitrified in liquid N_2_ in mother liquor supplemented with 30% [v/v] glycerol.

X-ray diffraction data were collected on beamline PXIII of the Swiss Light Source (SLS, Villigen, CH). Data processing used XDS/XSCALE^46^. sMolecular replacement was performed with Phaser^47^ using the TwcK/Stu structure as a search model. Refinement was as described above. Statistics for X-ray data processing and model refinement are given in **Table 1**.

### Kinase phosphotransfer assays

Kinase activity was monitored using the Kinase-Glo® Luminescent Kinase Assay (Promega) in white 96-well plates (Greiner bio-one) adapting reported protocols^45^. Here, 20µL of solution containing 5mg/mL BSA, 2.5mM MgCl_2_, 50mM Tris-HCl, 50mM NaCl, 0.5mM TCEP, pH 7.9 were incubated with the reaction partners to be investigated at final concentrations of 0.4mg/mL TwcK or NL-TwcK, 1µM ATP and 0.8mg/mL of one of the following peptides (commercially synthesized by ProteoGenix): kMLC_11-23_ (KKRARAATSNVFS^20,22^), kMLC ^T18P^ (KKRARAAT^P^SNVFS), kMLC ^S19P^ (KKRARAATS^P^NVFS), kMLC ^S23P^ (KKRARAATSNVFS^P^), kMLC (KKRARAATSNVF), kMLC ^T18P^ (KKRARAAT^P^SNVF), kMLC ^S19P^ (KKRARAATS^P^NVF) or MLC-4 (RQRPQRATSNVFA). Experiments in which one of the substrates was missing were adjusted to a final reaction volume of 50µL reaction buffer. This reaction mixture was incubated for 8min before 50µL of ATP-Glo-Kit® were added. The latter consumed the ATP remaining in the solution instantly generating a luminescence signal that was detected with a Infinite M200Pro luminometer (TECAN). The measured luminescence values were normalised to the negative control measurement of the inactive kinase mutant TwcK^K185A^ (previously reported^23^).

### Monitoring kinase phosphotransfer activity by ^31^P NMR spectroscopy

TwcK activity was monitored in real-time by applying ^31^P NMR spectroscopy. To start the reaction, 20µM MgCl_2_ was added to a solution containing 1mM ATP, 1mM kMLC_11-23_ and 10µM TwcK in 50mM Tris, 50mM NaCl, 0.5mM TCEP, pH 7.9 to a final concentration of 20µM. Immediately after the addition of MgCl_2_, the NMR sample tube was inserted into an Avance Neo 800MHz (18.8T) NMR spectrometer (Bruker) equipped with a QCI cryoprobe and a series of 28 1D ^31^P NMR spectra were recorded until the reaction was finished. Each spectrum was recorded using 608 scans and an inter-scan delay of 1.012s resulting in an experimental time of 20min, with a delay of 10min between each spectrum, at a constant temperature of 20.4°C. ^31^P NMR spectra were processed and analysed using Topspin 4.0.7 (Bruker). Product formation was quantified by integration of the resonance signal corresponding to kMLC_11-23_ phosphorylated at residue T18, emerging at a chemical shift of 3.16ppm over time, in the spectral range from 3.32 – 3.00 ppm. Data were plotted and further analysed using Origin 2019b (OriginLab).

1D ^31^P NMR reference spectra from pre-phosphorylated peptides (kMLC ^T18P^, kMLC ^S19P^, kMLC ^S23P^) were recorded using the same spectral parameters and buffer composition as described above. Each reference sample contained 1mM pre-phosphorylated peptide, 0.5mM ATP, 0.5mM ADP, 0.05mM AMP, 20µM MgCl_2_. 1D ^31^P NMR reference spectra acquired from nucleotides (AMP, ADP, ATP) were measured from samples containing 1mM nucleotide, 20µM MgCl_2_ with buffer composition as before.

### Identification of MLC-4 in the *C. elegans* genome

The amino acid sequence RAATSNVF, corresponding to the central segment of the kMLC peptide, was subjected to a similarity search against the non-redundant (nr) protein sequence database of *C. elegans* (taxid:6239) using Blast (https://blast.ncbi.nlm.nih.gov/Blast.cgi). Sequences were then filtered by imposing conservation of the TS residues as potential phosphorylation targets. The myosin regulatory light chain isoform 4 (MLC-4) (NCBI: NP_497700.1) emerged then as a candidate substrate for TwcK with the highest sequence similarity.

### CRISPR/Cas9 generation of *C. elegans* strain PHX9596 [*mlc-4(syb9596)*]

The *C. elegans* strain PHX9596 [*mlc-4(syb9596)*] that genomically expresses MLC-4 carrying C-terminally a 3xHA tag (MLC-4-HA) was generated by genome editing using CRISPR/Cas9. The procedure was performed commercially by SunyBiotech (www.sunybiotech.com) using the following materials (synonymous mutations are boxed):

PCR primers:

BG42-out-s: CTTTAGAGATGCTCCGATTA
BG42-out-a: ACATTTACAGCTTTGGCAAC

Sequencing primers:

BG42-oligo-a: TGAGTTTGCGAGCCAGTTTAAGCGT
BG42-out-a: ACATTTACAGCTTTGGCAAC

>BG42-syb9596
CTTTAGAGATGCTCCGATTAAGgtAtaatcgctgtaacttcggaacgaaaaaaagaagggaaacaggagtttctggc tcgaaaagcttgattttggacctcttaaaatcgtggaaaattcagagattttcgatttatcagccggaaaatttggt gtcgatttacgccggtcgtgtactcctagtgaagaagtttttgcttctttattttgcaaaaaattgcgataattttc gctttttattgtagattttgaaggaaaaaaatcgatgttgaaattacttgttttttcaacagaaaaaagtgcaaact tctccacggggagtacacgatctgcgtaaatcgacacaattttctataaaaaaaacacatatttttaccaaattttc caatgcaaaatagttttttttaaatcctaaaaccctaaaatatttattttttcctgaaaaataagccaattttcaat tttttttttctcagGGCGGACAATTCGACTACGTGGAGTTCACTCGTATGCTCAAACACGGAACC**AA** AAGGAT GAGGCTTACCCATACGATGTTCCAGATTACGCTGGTGGATCTTACCCATACGATGTTCCAGATTACGCTGGTGGATC TTACCCATACGATGTTCCAGATTACGCTTAAactggctcgcaaactcacacaaaattctcgagtttttcaagattca gttcgcttacacacgctttctttttcgcctcaaataattattatcattcatgctttttttcacaccatctatttatt gttgtattccacttcccccaataccatgtgcataaataaactcgaaaacggttgatttttattgtaatttaaagcta taatctttaaggattttttcacagattcgtttctcgggaagtttcaatcatcacaggagggtattgatgagctcgtc attcgtcggatttagaagcattatcagggataatggagatttttcgagttggcgggttatttctgcaaaaaattgaa cattttcaggattgtaatttccggcaaatcggcaaaccggcaatttgccgatttgccgggattttggccatagaaaa ttgcccaattttcgaaaaaaaaaagtttactaaccactgaaaatagtttgagtgttggcaagtgtacacttttgtga cttataaattgagcaaaatcggcacattgcagatgcatggagagcagaaaagaatggatagagtccttcgatttcgt tgccaaagctgtaaatgt

In the above, upper case represents coding sequence, and lower case is either intron, 3’UTR or downstream genomic sequence. The 2 nucleotides that are boxed were changed to prevent re-digestion by Cas9 but did not change the translated amino acids.

### Western blot demonstrating expression of MLC-4-HA in *mlc-4(syb9596)*

Total protein lysates from mixed-stage populations of wild type and *mlc-4(syb9596)* worms were prepared by the method of Hannak et al. (2002). Equal amounts of total protein were separated on a 12% polyacrylamide SDS Laemmli gel, transferred to nitrocellulose membrane, stained with Ponceau S, and reacted against anti HA antibodies (rabbit monoclonal cat. no. C29F4 from Cell Signaling Technology at 1:1,000 dilution), and then reacted with goat anti rabbit immunoglobulin G conjugated to HRP at 1:10,000 dilution and visualized by enhanced-chemiluminescence (ECL).

### Immunostaining of the pharynx and body wall muscle of *C. elegans*

To image strain EU573, *orEx2*[mlc-4p::mlc-4::GFP; rol-6(su1006)]^38^, that transgenically produces MLC-4-GFP, synchronized adult nematodes were immunostained and imaged as follows. To image both MLC-4-GFP and MHC C, ∼200 adult rollers were fixed and processed (as described^50^), and reacted with anti-GFP (rabbit polyclonal, Thermo Fisher cat. no. A11122) and anti-MHC C (mouse monoclonal 9.2.1 obtained from the University of Iowa Hybridoma Bank) each at 1:200 dilution. Secondary antibodies, also used at 1:200 dilution, were anti-rabbit Alexa 488 and anti-mouse Alexa 594 (Invitrogen).

PHX665 is a CRISPR strain in which mCitrine was placed N-terminally to the kinase domain of twitchin^31^. To image both twitchin-mCitrine and MHC C from strain PHX665 [*unc-22(syb665)*], we fixed and processed adults as described^50^. MHC C was imaged as described above, whereas twitchin-mCitrine was imaged by direct fluorescence using a bandpass filter at 505-550 nm.

To study strain PHX9596 [*mlc-4(syb9596)*], adults were fixed (method in ^50^) and reacted with anti-HA (rabbit monoclonal cat. no. C29F4 from Cell Signaling Technology at 1:200 dilution) and anti-MHC B (mouse monoclonal 5-8 obtained from the University of Iowa Hybridoma Bank) at 1:200 dilution, to image body wall muscle, and anti-HA and anti-MHC C to image the pharyngeal muscle. Secondary antibodies, also used at 1:200 dilution, were anti-rabbit Alexa 488 and anti-mouse Alexa 594 (Invitrogen).

In all instances, images were captured at room temperature with a Zeiss confocal system (LSM510) equipped with an Axiovert 100M microscope and an Apochromat X63/1.4 numerical aperture oil immersion objective in 1X mode. Colour balances were adjusted by using Adobe Photoshop (Adobe, San Jose, CA).

## Data Availability

Model coordinates and structure factors have been deposited with the Protein Data Bank: TwcK/Stu, PDB entry 29QN; TwcK/Stu/kMLC_13-22_, PDB entry 29UJ. X-ray diffraction images for TwcK/Stu and TwcK/Stu/kMLC_13-22_ have been deposited with Zenodo. TwcK/Stu corresponds to entries 10.5281/zenodo.11198709, 10.5281/zenodo.11203533, 10.5281/zenodo.11203615, 10.5281/zenodo.11203671. TwcK/Stu/kMLC_13-22_ corresponds to entry 10.5281/zenodo.18599684.

## Acknowledgments

We thank the Swiss Light Source (SLS) for synchrotron radiation time. We acknowledge the financial support of the Deutsche Forschungsgemeinschaft (DFG, German Research Foundation) SFB1756 – 550938463 to OM and MK and grant 2050009 from the National Science Foundation to GMB. We thank the University of Konstanz for permanent investment on NMR spectroscopic infrastructure.

## Authors contributions

PG, RW, TZ, and OM undertook sample preparation, crystallographic studies and luminescence activity studies; PG, TD and OM performed bioinformatics analyses; FB and MK implemented NMR spectroscopy and all its related analyses; HQ and GB contributed all *C. elegans* related work; O.M. conceived the study; PG and OM wrote the paper with the contribution of all authors.

## SUPPLEMENTARY MATERIALS

**Fig S1:**
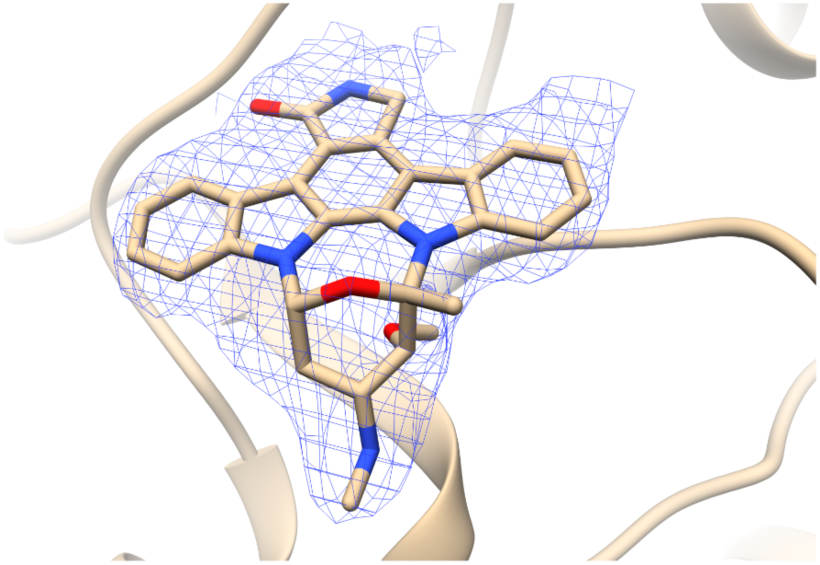
TwcK ligands. (2mF_obs_-DF_calc_)α_calc_ electron density maps contoured at 1σ of staurosporine bound into the ATP-binding pocket.

**Fig S2:**
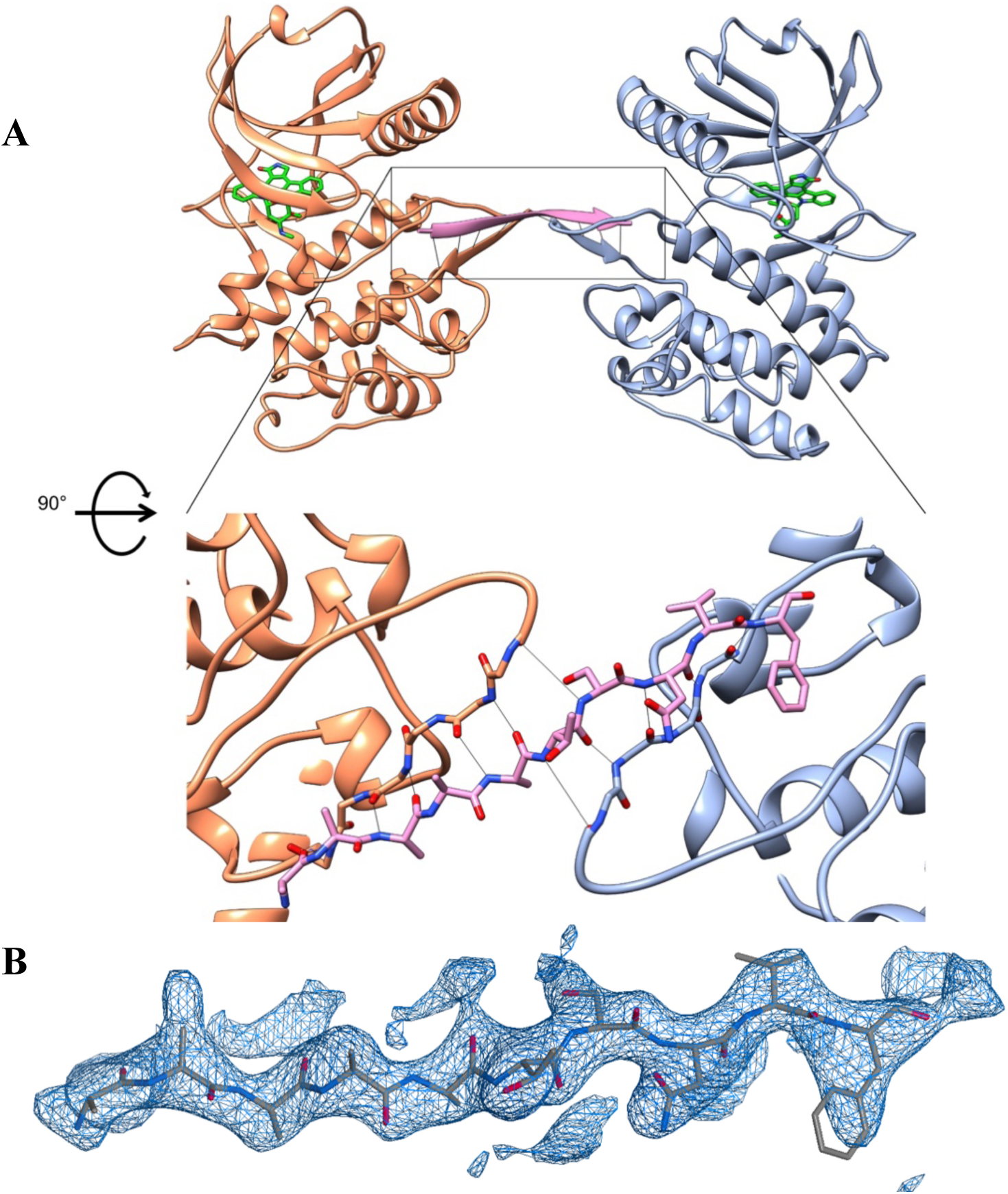
kMLC_13-22_ peptide binding by TwcK. **A.** Composition of the asymmetric unit of TwcK/Stu/kMLC_13-22_ crystals, consisting of two molecular copies of TwcK (orange, blue) and a kMLC_13-22_ peptide (pink). The interaction of the peptide with TwcK is mediated by main chain groups, with no involvement of lateral groups, which are poorly assignable in electron density maps; **B.** (2mF_obs_-DF_calc_)α_calc_ electron density maps contoured at 1σ of kMLC_13-22_.

**Fig S3:**
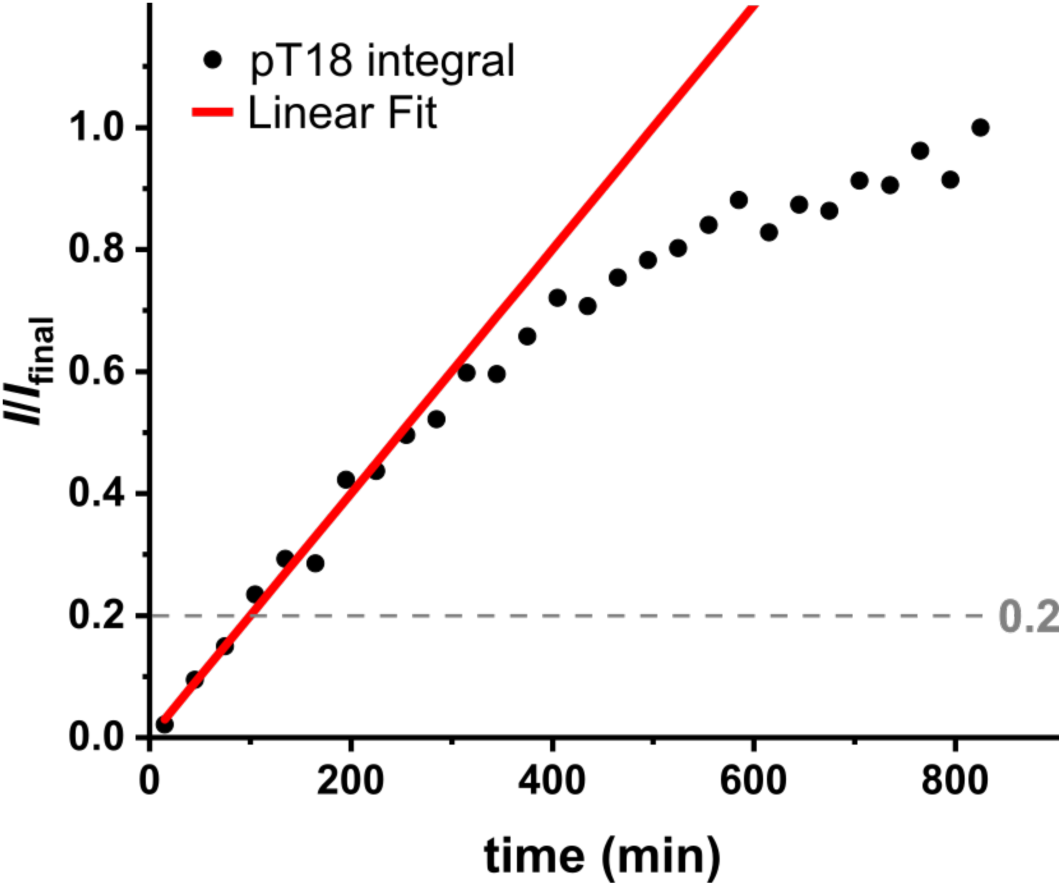
Turnover number of TwcK on the kMLC_11-23_ peptide substrate. To illustrate the reaction course, recorded integrals (*I*) were normalised to the final integral (*I*_final_) of the emerging resonance signal corresponding to the phosphorylated peptide. The enzymatic reaction investigated is a bi-substrate reaction (kMLC_11-23_ + ATP ⇌ p-kMLC_11-23_ +ADP). At the beginning of the reaction, the substrate concentrations (*c*[ATP] = 1mM, *c*[kMLC_11-23_] = 1mM) are much higher than the enzyme concentration (*c*[TwcK] = 10µM) and saturating conditions are assumed, so that the reaction is initially of zero order with the concentration change of the reactants varying linearly with time. As the reaction takes place and substrates are consumed, the reaction velocity decreases according to the Michaelis-Menten theory. A linear function was fitted to the integrals normalised to the final integral during the initial reaction time (*I*/*I*_final_ < 0.2) leading to determine the rate constant of the increase, *k,* as *k* = (2.00 ± 0.08) * 10^−3^ min^−1^. This parameter was then converted to the catalytic turnover number *k*_cat_ by multiplying by the assumed final concentration (1 mM) and normalized to the enzyme concentration (10 µM). The turnover number determined in this way was *k*_cat_ = (3.34 ± 0.14) * 10^−3^ s^−1^.

**Fig S4:**
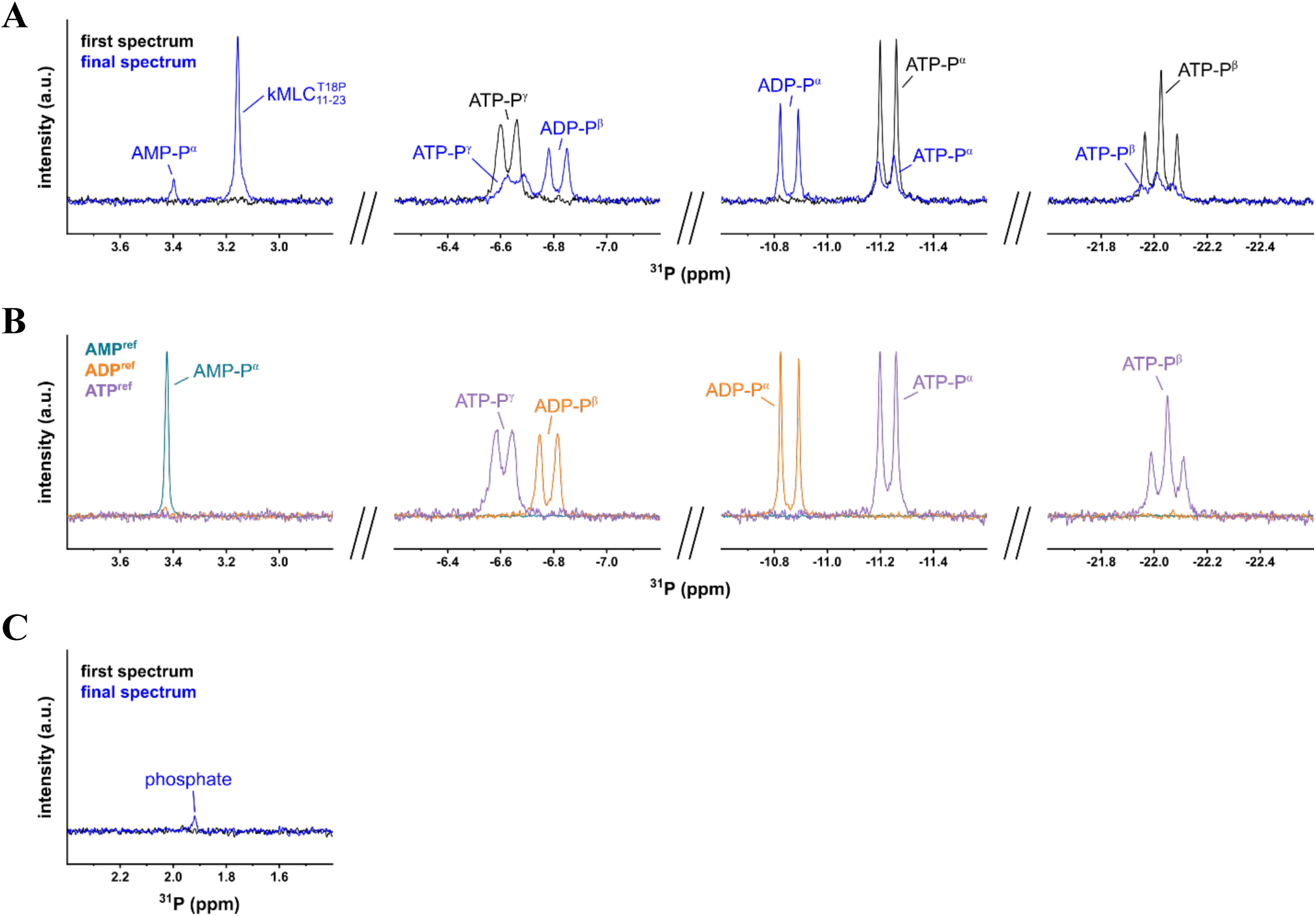
1D ^31^P NMR reference spectra. Spectral ranges (3.8 – 2.8 ppm, −6.2 – −7.2 ppm, −10.6 – −11.6 ppm, −21.6 – −22.6 ppm) of ^31^P NMR spectra recorded on the reaction mixture and reference spectra of ATP, ADP and AMP are shown. **A.** The “first spectrum” (black) shows the spectrum of a reaction mixture containing TwcK, ATP and kMLC_11-23_ before addition of MgCl_2_, the “final spectrum” (blue) represents the spectrum of TwcK, ATP, MgCl_2_ and kMLC_11-23_ after 825 min incubation; **B.** ATP (purple), ADP (orange), AMP (teal) reference spectra contain the respective nucleotides and MgCl_2_. **C** Spectral range (2.4 – 1.4 ppm) showing the resonance signal corresponding to free phosphate at 1.92 ppm. The emerging resonance signals at −6.81 ppm and −10.86 ppm in the reaction mixture are assigned to ADP that is produced during the reaction. ADP decomposes over time into free phosphate (1.92 ppm) and AMP (3.40 ppm). An experimental mixture consisting of ATP, ADP, AMP and free phosphate leads to minor deviations of the chemical shift compared to individual reference spectra.

**Fig S5:**
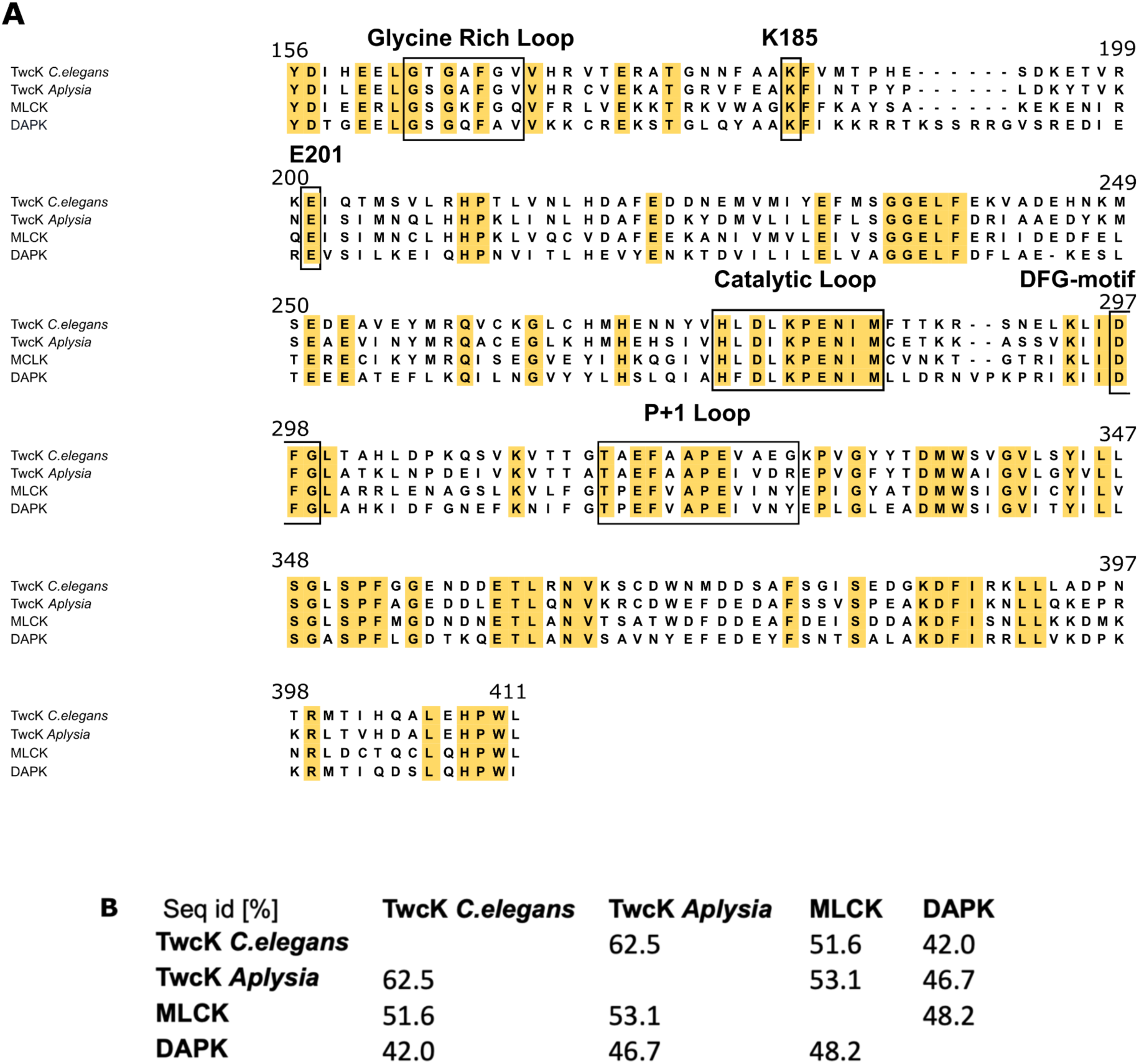
Multiple Sequence Alignment of DMT kinases that perform phosphotransfer activity on myosin light chain (MLC) proteins. **A.** Multiple sequence alignment of *C. elegans* TwcK (Uniprot Q23551, residues 5538-5793; PDB 3UTO residues 156-411), *Aplysia* TwcK (Uniprot Q16980 residues 47-302), MLCK (Uniprot Q15746 residues 1464-1719), DAPK (Uniprot P53355 residues 13-275). Sequence identity is highlighted in yellow and catalytic relevant motifs are boxed. **B.** Sequence identity matrix of the four kinases aligned in A.

